# Secondary structure of subgenomic RNA M of SARS-CoV-2

**DOI:** 10.1101/2021.10.11.463917

**Authors:** Marta Soszynska-Jozwiak, Ryszard Kierzek, Elzbieta Kierzek

## Abstract

SARS-CoV-2 belongs the *Coronavirinae* family. As other coronaviruses, SARS-CoV-2 is enveloped and possesses positive-sense, single-stranded RNA genome of ∼ 30 kb. Genome RNA is used as the template for replication and transcription. During these processes, positive-sense genomic RNA (gRNA) and subgenomic RNAs (sgRNAs) are created. Several studies showed importance of genomic RNA secondary structure in SARS-CoV-2 replication. However, the structure of sgRNAs have remained largely unsolved so far. In this study, we performed probing of sgRNA M of SARS-CoV-2 *in vitro*. This is the first experimentally informed secondary structure model of sgRNA M, which presents features likely to be important in sgRNA M function. The knowledge about sgRNA M provides insights to better understand virus biology and could be used for designing new therapeutics.

## Introduction

Severe acute respiratory syndrome coronavirus 2 (SARS-CoV-2) causes Coronavirus disease 19 (COVID-19) and is responsible for infecting a large number of people all over the word, resulting in death of many people, especially among older. Moreover, SARS-CoV-2 causes disruptions to health service, travel, trade, education as well as has a negative impact on people’s physical and mental health [1]. SARS-CoV-2 belongs to the Betacoronavirus genus and is a member of the *Coronavirinae* family which also includes alpha-, gamma- and deltacoronaviruses [2]. As other coronaviruses, SARS-CoV-2 is enveloped, and possess three structural proteins: membrane protein (M), spike protein (S), and envelope protein (E), while nucleocapsid protein (N) protects viral RNA genome by forming a capsid [3]. SARS-CoV-2 also contains sixteen non-structural proteins (nsp1−16) and accessory proteins [4], [5]. SARS-CoV-2 represent RNA viruses and possess positive-sense, single-stranded RNA genome of ∼ 30 kb [6]. The genomic RNA has a 5′ cap and 3′ polyA tail and is used for translation to produce two large overlapping polyproteins (pp): pp1a and pp1ab. Polyproteins pp1a and pp1ab contain the nsps 1–11 and 1–16, respectively. Many of them with N protein create replicase-transcriptase complex (RTC) [7]. Genome RNA is used as the template for replication and transcription by RTC. These processes result in generation negative-sense RNA intermediates, that serve as temple for production positive-sense genomic RNA (gRNA) and subgenomic RNAs (sgRNAs). The gRNA is packaged to progeny virions or is used for translation, while sgRNAs encode conserved structural proteins, nucleocapsid protein and several accessory proteins [8]–[11].

Each sgRNA possesses a short 5’-terminal leader sequence derived from the 5’ end of the genome. Transcription regulatory sequences (TRS) is necessary to add leader sequence to sgRNA [12]. TRS is located at the 3’ end of the leader sequence (leader TRS, TRS-L), as well as TRSs are located upstream of the genes in the 3’-proximal part of the genome (body TRSs, TRS-B). TRSs contain a conserved 6–7 nt core sequence (CS) surrounded by variable sequences. During negative-strand synthesis, RNA*-*dependent RNA polymerase (RdRP) pauses when it crosses a body TRS and switches the template to leader TRS. As a result sgRNA possess a leader sequence derived from the 5′ untranslated region of the genome and a TRS 5′ of the open reading frame [13], [14].

RNA secondary structure of untranslated and coding regions play a key role in viral replication cycle [15]. Secondary structure of RNA of SARS-CoV-2 has been intensively studied. Bioinformatic analysis showed that SARS-CoV-2 genome has almost twice the propensity to form secondary structure than one of the most structured RNA genomes in nature [16]. Recently, detailed secondary structure models for the extended 5’ UTR, frameshifting stimulation element, 3’ UTR and regions of the SARS-CoV-2 viral genome that have low propensity for RNA secondary structure and are conserved within SARS-CoV-2 strains were predicted [17] [18]. SARS-CoV-2 was also investigated for the presence of large-scale internal RNA base pairing (genome-scale ordered RNA structure) (GORS)) in its genome. This analysis showed existence of 657 stem-loops structures and 2015 duplexes [19]. Data revealed that regions containing the highest amount of structure of SARS-CoV-2 genome are the 5’ end as well as regions corresponding to glycoproteins spike S and membrane M [20]. Lately, RNA structure probing of the full SARS-CoV-2 coronavirus genome both *in vitro* and in living infected cells was published [21]–[24]. Recently, using RNA structure probing with Nanopore direct-RNA sequencing, NAI reactivity of 3a, E, M, 6, 7a, 7b, 8 and N sgRNAs were measured [25]. However, the structure model of sgRNA M has not been determined and discussed so far.

In this study, we performed probing of the model of sgRNA M of SARS-CoV-2 *in vitro*. We use SHAPE method and chemical mapping with DMS and CMCT to obtain secondary structure of sgRNA M. This is the first experimentally informed secondary structure model of sgRNA M, which presents features likely to be important in sgRNA M function. A full set of chemical reagents allow to predict high accuracy structure. A comprehensive understanding of sgRNA M structure will provide important insights to better understand virus biology and could be used for designing new therapeutics.

## Results and Discussion

### Secondary structure of sgRNA M of SARS-CoV-2

Model of subgenomic RNA M of SARS-CoV-2 (sgRNA M) was obtained by adding leader sequence to M coding sequence from the SARS-CoV-2 strain Slovakia/SK-BMC5/2020 using PCR reactions. This process allowed achieving RNA with “leader to body fusion” that is characteristic for sgRNA of Coranaviridae family and taking place during transcription [14]. Chemical mapping was used to determine secondary structure of sgRNA M. *In vitro* transcribed sgRNA M was folding in folding buffer (300mM NaCl, 5mM MgCl_2_, 50mM HEPES, pH 7.5) to obtain single RNA conformation, as assessed by non-denaturing agarose gel. Chemical mapping was performed at 37°C with DMS (methylates N1 of A and N3 of C when unpaired), CMCT (modifies N3 of U and N1 of G when unpaired) and NMIA (SHAPE method, modified flexible 2’-hydroxyl of ribose) [26][27][28]. The data from chemical mapping were analyzed by reverse transcription followed by capillary electrophoresis.

DMS modified strongly 123 nucleotides (nt), meaning that 32.6% of all adenosines and cytidines in sgRNA M were structurally accessible. CMCT modified 106 nts, which represents 40.9% of all uridines. The SHAPE reagent NMIA modified strongly 119 nts and 56 nts undergo moderate modifications, that is together 22.2% of sgRNA M nucleotides. Chemical mapping data and SHAPE data are complementary to each other and show eight the most flexible regions (each containing at least 7 flexible nucleotides in sequence): 36-43, 76-86, 260-267, 469-481, 487-494, 596-603, 612-621, 729-743 (Figure 1).

**Figure 1.**
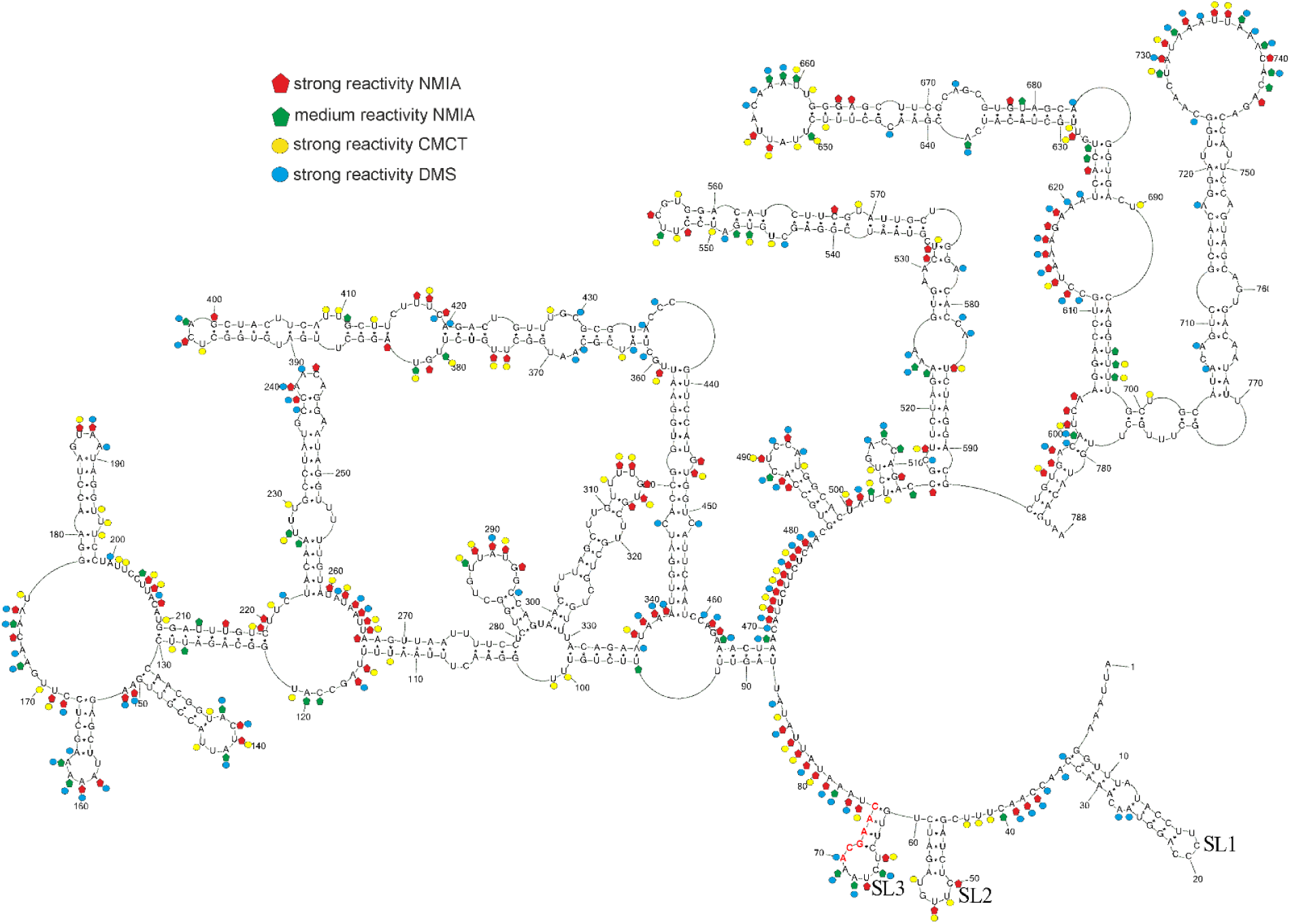
sgRNA M model predicted by RNAstructure 6.2 using experimental data as constraints. Strong DMS and CMCT modifications, as well as SHAPE reactivities converted to pseudo-free energies were used. The numbering of sgRNA M is from its 5′ end. The AUG start codon spans nucleotides 120–122. Red nucleotides indicate TRS sequences. Hairpins SL1, SL2 and SL3 are indicated.

### Base pair probabilities

To assess prediction quality and identify well-defined structural regions we calculated the secondary structure partition function using RNAstructure 6.2 and, from this, determined the base pair probabilities for model pairs [29]. For the partition function calculations, experimental data were included. Results indicate that there are several regions with paired and unpaired nucleotides of more than 90% probability: 1-121, 131-210, 220-502, 707-766. Additionally, all single stranded regions are well defined by having low probability of pairing (Figure 2).

**Figure 2.**
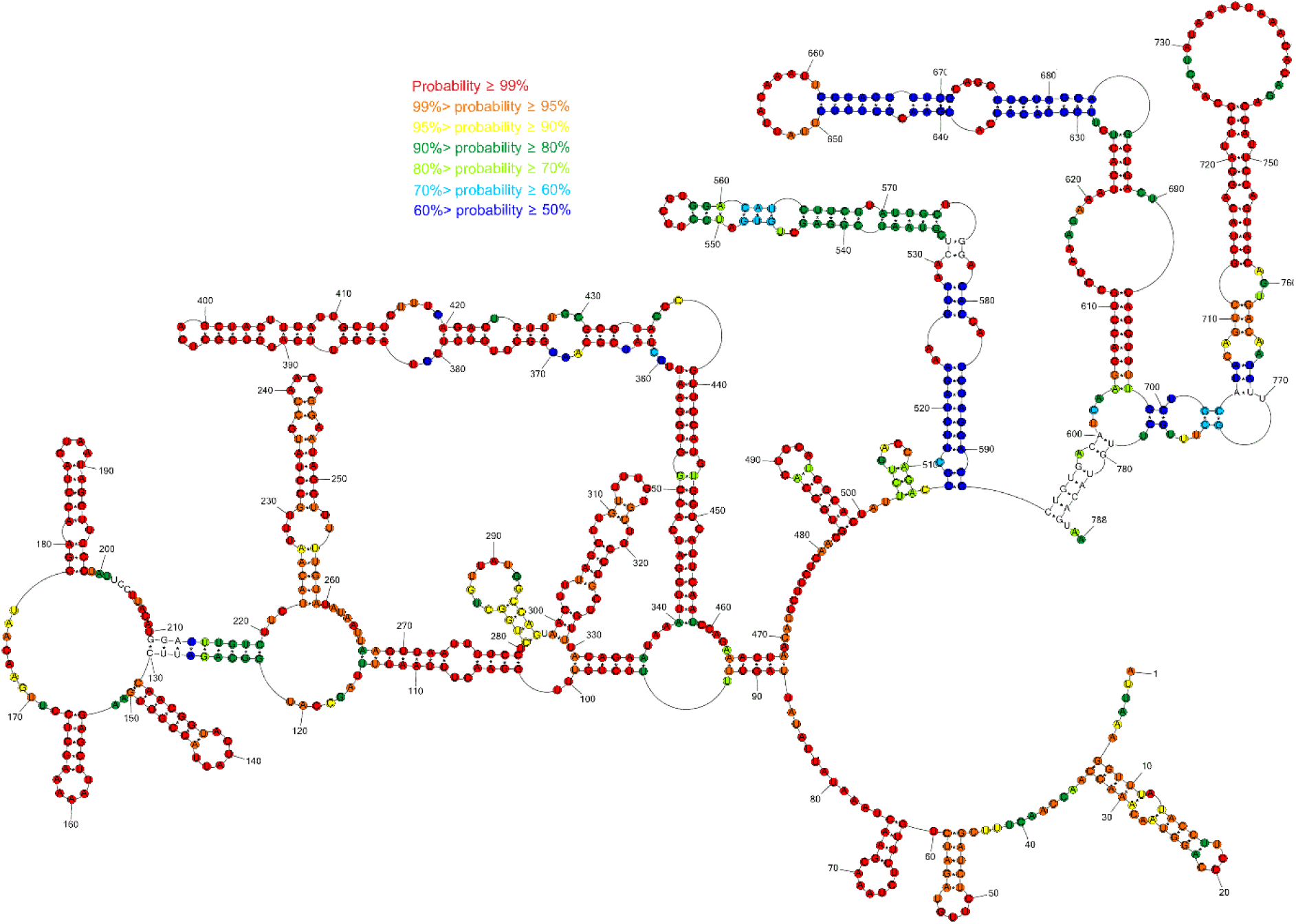
Predicted probability of nucleotides being paired or single stranded in sgRNA M using RNAstructure program. Probability lower than 50% is not colored. The partition function calculation incorporated restraints from strong reactivity of DMS and CMCT as well as SHAPE reactivities converted to pseudo-energies.

### Model of secondary structure of sgRNA M

To predict the secondary structure of sgRNA M based on the experimental probing data, the results of chemical mapping were used to constrain predictions in the RNAstructure 6.2 program. SHAPE data were loaded as pseudoenergy constraints (the energy contribution of SHAPE reactive nt were penalized) and DMS, and CMCT modifications were included as chemical mapping constraints (reactive nt are forbidden to be in Watson-Crick base pairs flanked by Watson-Crick base pairs).

Model of sgRNA M is highly structured with plenty of accessible bulges and loops (Figure 1). RNA motifs of sgRNA M model are thermodynamically stable (dG=-418,5 kcal/mol) and have high calculated base pair probability. We showed that most of the inaccessible regions by chemical mapping, correspond to areas containing base pairs. 5’end of the sgRNA M model was folded in three hairpins: SL1, SL2, SL3. This three hairpins also occurs in 5’UTR of SARS-CoV-2 in its 5’ 300 nt fragment [21], [30], [31], in *in vitro* model of whole genome and also in-cell model [21]–[24], [32]. These hairpins are also in good agreement with structural–phylogenetic analysis of group IIb coronaviruses [4] and *in silico* prediction of whole SARS-CoV-2 genome [17], [18]. Moreover, folding of a 5’ leader sequence of sgRNA M is in agreement with study of secondary structure of subgenomic RNA N [16]. This investigation showed that 5’ leader sequence folds almost autonomously in the sgRNA N, with the exception of a few poorly determined long-range interactions [16]. SL1 is the most variable among SARS-CoV-2 variants [30], generally possessing mismatches, bulges and a high number of A–U and U–A base pairs. This fact causes less thermodynamically stability of SL1 than SL2 and SL3. On the other hand, this feature is important for the replication of Mouse hepatitis virus (MHV), well-studied members of coronaviridae family [33]. SL2 is conserved in all CoVs, typically containing a pentaloop stacked on a 5 base-pair stem and creating U-turn motif. This hairpin plays a critical role in MHV replication and translation [34]. SL3 is conserved only in subgroups of beta and gammaCoV [4] and contains TRS-L sequences, that take part in discontinuous transcription [4], [13].

Recently, prediction of interaction between SARS-CoV-2 genome and human proteome indicated that highly structured region at the 5′ end had the large number of interactions with proteins such as 1) ATP-dependent RNA helicase - DDX1, that was previously reported to be essential for Avian infectious bronchitis coronavirus replication [35] and 2) ADAR double-stranded RNA-specific editases, that catalyse the hydrolytic deamination of adenosine to inosine, result in affecting viral protein synthesis, proliferation and infectivity [36], and 3) 2′-5′-oligoadenylate synthetases that control viral RNA degradation [20], [37], [38]. Some of these proteins could interact with a leader sequence of sgRNA. This fact confirmed experiments with DDX1 knockdown that reduced the number of sgRNA in SARS-CoV-1 infected cells [39]. These finding and preservation of 5’UTR motifs in sgRNA M could indicate the similar interactions with sgRNA M. Moreover, interactions between SARS-CoV-2 genome as well as sgRNAs and host RNAs were revealed. However, SAR-CoV-2 genome and sgRNAs take part in different interactions with host RNAs [25], [32].

Generally, presented model of secondary structure of sgRNA M and its corresponding region of the whole SARS-CoV-2 genome obtained by probing *in vivo* [22] are different (Figure 3). It is in agreement with previous study about *in vivo* RNA-RNA interactome of the full-length SARS-CoV-2 genome and several subgenomic mRNAs. This investigation revealed that the viral genome and subgenomes adopt alternative topologies inside cell [32]. However, some regions in the presented model of sgRNA M fold in the same structure as in genome M region in infected cells. Motifs: 153-167, 178-198, 223-259, 299-329, 340-351/447-458, 362-435, 484-498, 514-592, 604-698, 712-759 are consistent with in-cell secondary structure model of SARS-CoV-2 genome [22]. However, hairpin 514-592 is longer about 3 base pairs and possesses an additionally internal loop in-cell model. In turn, hairpins 131-150 and 281-297 are almost identical to that of corresponding region of SARS-CoV-2 genome that was mapped in cells [22]. In our model, loops are longer than corresponding region in-cell model. On the other hand, small motif 503-512 is only hairpin-structure characteristic for sgRNA M, and does not exist in SARS-CoV-2 genome. Determined sgRNA M secondary structure is also similar to corresponding regions of other published whole genome of SARS-CoV-2 models [23, 24]. This similarity between sgRNA M structure and in-cell determined structure of fragment M in whole genome context is surprising due to the fact that RNA structure *in vitro* and *in vivo* could be significantly different. *In vivo* interaction between RNA and proteins or other molecules can influence secondary structure [40]. This data indicated that the sequence and thermodynamics alone are major determinants of sgRNA M motifs formation. All these results point to these stable motifs being functionally significant. Furthermore, experiments and computational analysis have shown that large amounts of double-stranded regions have a strong propensity to interact with proteins and act as scaffolds for RNA-binding proteins [41]–[43]. sgRNA M is very structured and it is possible for stable helices to interplay with proteins.

**Figure 3.**
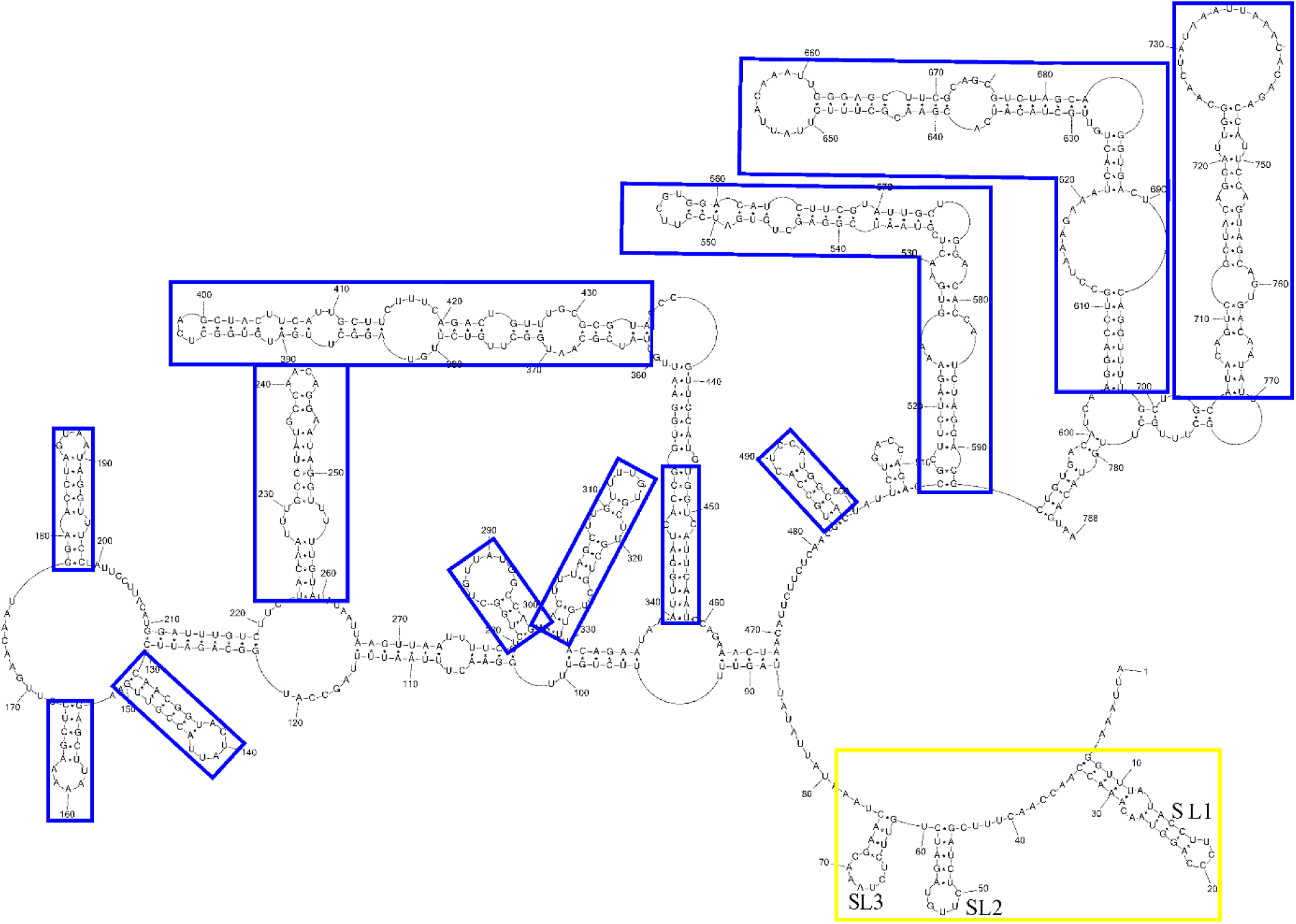
Comparison of secondary structure of sgRNA M and its corresponding region of SARS-CoV-2 genome mapped in-cells. Blue rectangle indicates the same base pairs within sgRNA M model and corresponding region of SARS-CoV-2 genome mapped in-cells [22]. Yellow rectangle indicates motifs of leader sequence.

### Local structural motifs in sgRNA M are mostly independent of leader sequence

We compared our sgRNA M model with the corresponding region of *in vitro* model of whole SARS-CoV-2 genome [21] to check the influence of 5’leader sequence on folding of subgenomic RNA. We indicated that the structures are different. This feature is consistent with study with subgenomic RNA N and its corresponding region in SARS-CoV-2 genome. This data indicated that the same RNA sequences can fold in different structures in the subgenomic and genomic contexts [16]. However, some local motifs (131-150, 153-167, 181-194, 223-259, 267-279/102-114, 280-297, 299-329, 331-335/101-105, 340-358/439-458, 372-426, 483-499, 514-592, 604-698, 712-758 within the subgenomic RNA and SARS-CoV-2 genome are identical. Motifs 604-698 is slightly different in both models. Similar motifs are stable independently of the neighbouring regions (leader sequence in sgRNA M and sequences upstream/downstream of the M region in SARS-CoV-2 genome). It is possible that the existence and appropriate folding of neighbouring region has some, but relatively small, influence on some local motifs of sgRNA M. Some local structure motifs are identical within *in vitro* and *in vivo* SARS-CoV-2 genome models [21].

## Conclusion

The presented herein for the first time experimentally driven secondary structure of sgRNA M contains unique features likely to be important for its functions. The structure includes also several the same distinct motifs as genomic M fragment of SARS-CoV-2 genome. The new knowledge about sgRNA M provides insights to better understand virus biology and could be used for anti-SARS-CoV-2 strategies and designing new therapeutics.

## Material and Method

### Experimental Constructs

DNA temples for synthesis of sgRNA M was obtained in several steps. Firstly, reverse transcription was carried out using SuperScript III (Thermo Fisher Scientific) with Random Primers Mix (New England BioLabs) on RNA of SARS-CoV-2 from stain Slovakia/SK-BMC5/2020 (received from https://www.european-virus-archive.com). Next, three PCR reactions using cDNA as template and specific primers (F1 and RM, F2 and RM, F3 and RM, Table) were done to add leader sequence and transcription promoter on 5’ end of sgRNA M. After this steps, primers FC, RC were used to add EcoRI site on 5’ end and Pst I site on 3’ end of template of M. DNA was purified using Pure Link ™ PCR Micro Kit (Thermo Fisher Scientific). The DNA template was cloned into pUC19 and sequenced using the M13F, M13R for confirmation of proper sequence (Table).

### Oligonucleotides synthesis and labelling

Primers for reverse transcription were synthesized by the phosphoramidite approach on MerMade synthesizer. Primers for reverse transcription were synthesized with fluorophores: 5-FAM and 6-JOE on 5’-end (Table 2).Primers were deprotected and purified according to published protocols [44] [45]. Concentrations of all oligonucleotides were measured using a Spectrophotometer UV (NanoDrop2000 Thermo Scientific). Primers for PCR were purchase from Sigma-Aldrich.

### RNA synthesis

DNA template for *in vitro* transcriptions of sgRNA M was obtained by PCR from plasmid puC19 using primers FM, RM (Table). DNA was purified using Pure Link ™ PCR Micro Kit (Thermo Fisher Scientific). *In vitro* transcription reaction was performed using a MEGAscript™ T7 Transcription Kit (Thermo Fisher Scientific) according to manufacture protocol. Product was purified using RNeasy MiniElute Cleanup Kit (Qiagen). Integrity and purity of samples were checked on agarose gel.

### RNA folding

Before each experiment RNA was folded in the same manner. RNA was heated to 80°C in water for 5 min. and slowly cooled to 50°C. At this temperature 2-times concentrated folding buffer was added and samples were slowly cooled to 37°C. Final buffer contained 300 mM NaCl, 5 mM MgCl_2_, 50 mM HEPES, pH 7.5. Folding was analysed by native gel electrophoresis using 0.8% agarose gel running at 4 °C with low voltage. Under these conditions, one band was observed (Supporting_Information_2).

### Chemical mapping using NMIA, DMS and CMCT

Folding of RNA was carried out as described above. Next, chemical mapping was conducted according to published procedures with appropriate optimizations [26] [27] [28]. Briefly, 5.6 mM of NMIA (N-methylisatoic anhydride) or 30 mM of CMCT (1-cyclohexyl-(2-morpholinoethyl) carbodiimide metho-p-toluene sulfonate) or 0.18% of DMS (dimethyl sulfate) were used in mapping reactions. Chemical mapping was performed at 37°C with DMS, CMCT or NMIA for 15, 30 or 40 min, respectively. Parallel, control reactions were done in the same condition but without mapping reagents. Modified nucleotides were read-out by primer extension using a stoichiometry of 2 pmol primer/2 pmol RNA. Primer extension was performed at 55°C with reverse transcriptase SuperScript III (Thermo Fisher Scientific) using manufacturer’s protocol. Next, cDNA fragments and ddNTP ladders were separated by capillary electrophoresis (Laboratory of Molecular Biology Techniques at Adam Mickiewicz University in Poznan). Primers labeled with 6-FAM were used for detection of modification by DMS, CMCT or NMIA and control reaction without mapping reagents and resolved in two capillaries (reaction and control) with ddNTP ladders. Primers labeled with 5-JOE were used for ddNTP ladders (most often ddATP). The experiments were performed in at least technical triplicate with the average results presented. For obtained reactivity of each nucleotide the standard deviation (SD) is calculated (Supporting_Information_1).

### Processing of chemical mapping data

QuShape program was used to analyze mapping data according to published method [46]. NMIA reactivities was normalized by QuShape program using model-free statistics to a scale spanning 0 to ∼2, where zero indicates no reactivity and 1.0 is the lowaverage intensity for highly reactive RNA positions [46]. Nucleotides with normalized SHAPE reactivities 0–0.5, 0.5–0.7, and >0.7 correspond to unreactive, moderately reactive, and highly reactive positions, respectively. Nucleotides with no data were designated as −999. Normalized SHAPE reactivities from extension reaction of each primer were processed independently. DMS and CMCT modifications analysis was conducted similar to NMIA reactivity calculations, except that only strong modification were used in RNAstructure prediction.

Chemical mapping results were used in the RNAstructure [47] for prediction of secondary structure of sgRNA M. Normalized SHAPE reactivity (as described above) was used in RNAstructure 6.2 through “Read SHAPE reactivity—pseudo free energy” mode with slope 1.8 and intercept −0.6kcal/mol [48]. DMS and CMCT strong reactivities were introduced in the same prediction using “chemical modification” mode [49].

**Table 1.**
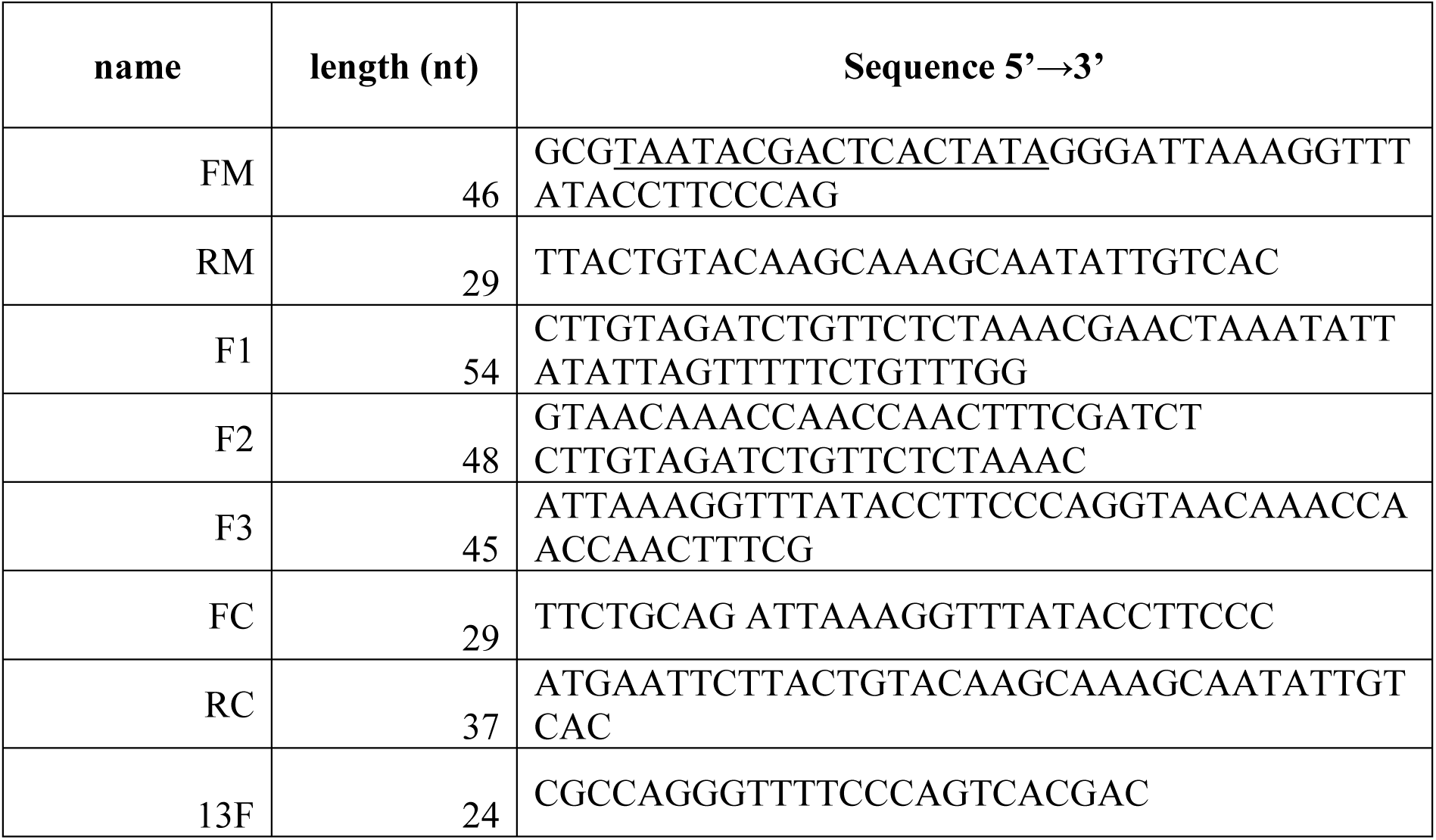
Primers for polymerase chain reaction using to obtain DNA template for sgRNA M. The underlined nucleotides residues are the polymerase T7 promoter.

**Table 1.**
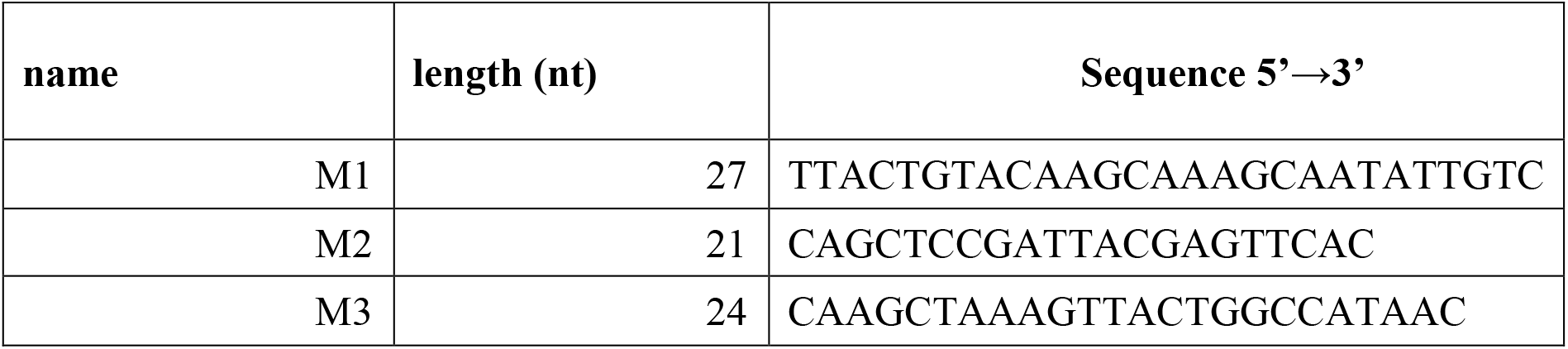
Primers for reverse transcription. Each primer was labeled with 5-FAM or 6-JOE at 5’ end.

## Supporting information

Suporting Information_2

