## Supplementary material for "Secondary structure of subgenomic RNA M of SARS-CoV-2": Suporting Information_2

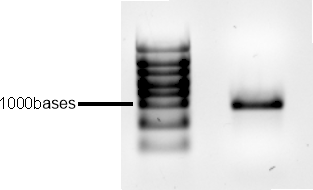

Figure S1. SgRNA M after folding by heating in 80°C for 5 min. in water and slowly cooled to 50°C. In 50°C buffer was added and slowly cooled to 37°C. Folding buffer: 300mM NaCl, 5mM MgCl_2_, 50mM HEPES, pH 7.5.

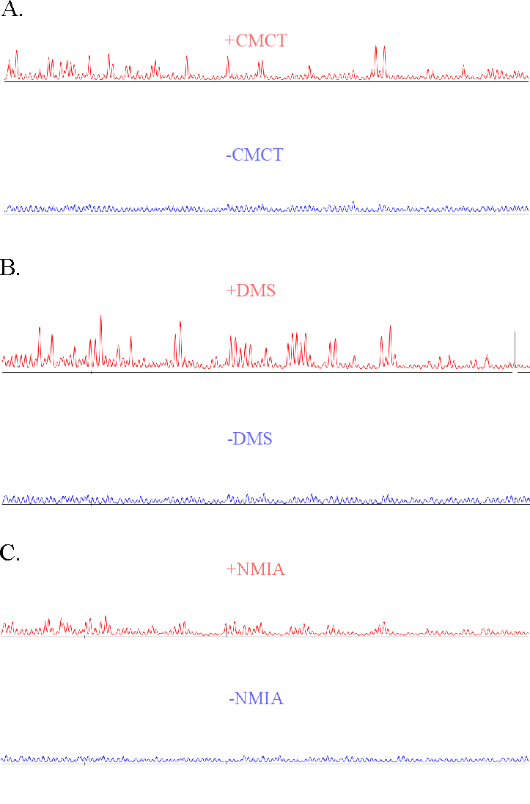

Figure S2. A-Example of capillary electrophoresis raw data for 231-107 nt. showing CMCT modified RNA (red line), unmodified control (blue line), B- Example of capillary electrophoresis raw data for 231-107 nt. showing DMS modified RNA (red line), unmodified control (blue line), C-Example of capillary electrophoresis raw data for 231-107 nt. showing NMIA modified RNA (red line), unmodified control (blue line).

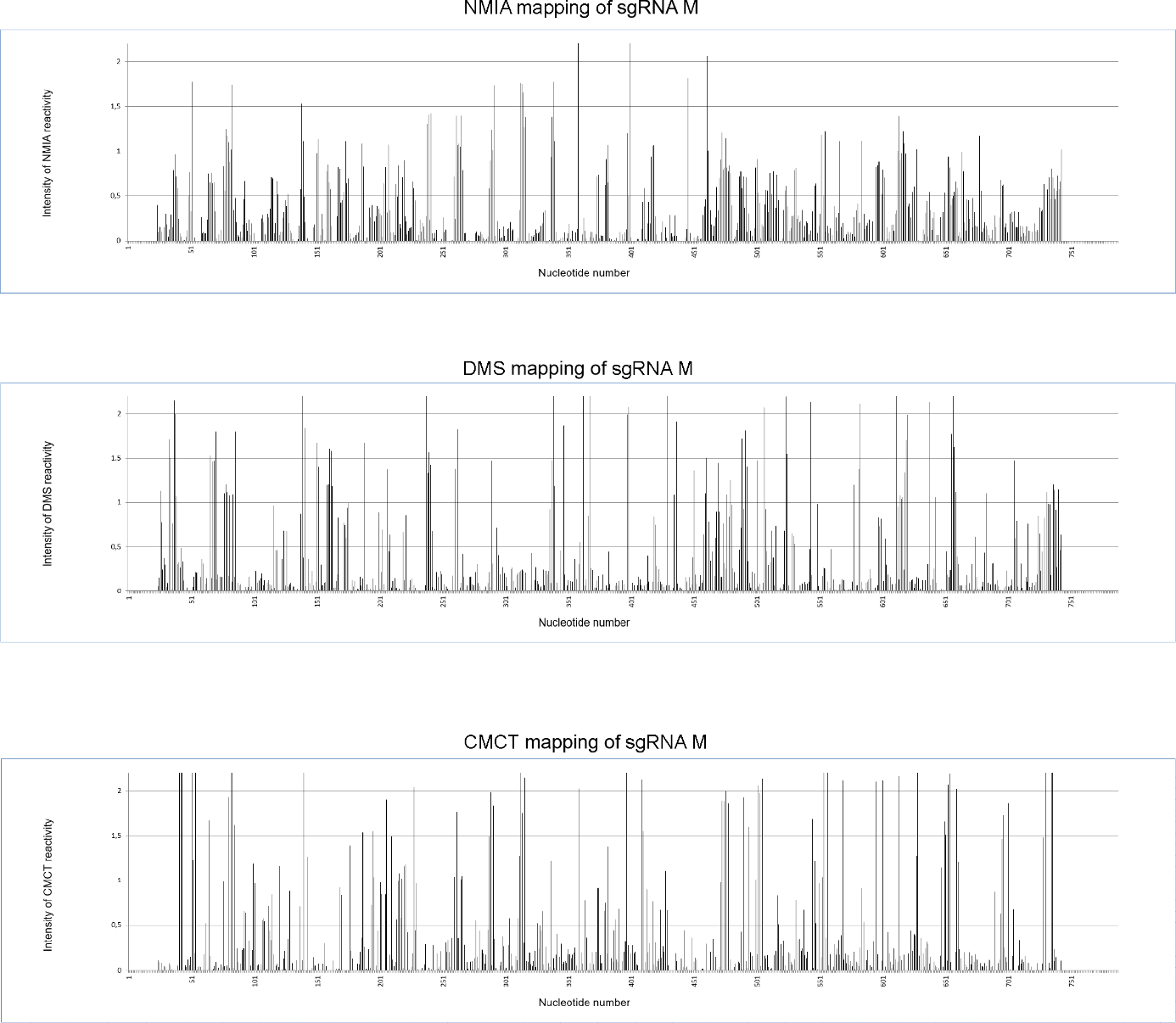

Figure S3. sgRNA M nucleotides reactivity diagrams. The sgRNA M chemical mapping experiments were performed at 37 °C with NMIA, DMS and CMCT.
